# The *Chromobacterium* Volatilome is Strongly Influenced by Growth on Liquid versus Solid Media

**DOI:** 10.64898/2026.03.19.712466

**Authors:** Jake A. Drewes, Jenna Diefenderfer, Diana Ramirez, Trenton J. Davis, Emily A. Higgins Keppler, Scott D. Soby, Heather D. Bean

## Abstract

The study of microbial volatile organic compounds (mVOCs) is a growing area of research, with applications ranging from agriculture to human health. The majority of the mVOC data are from *in vitro* liquid cultures, while few analyses of bacterial and fungal volatilomes on solid media cultures exist. Studies comparing liquid versus solid cultures of bacteria and fungi show significant changes to the soluble metabolites that are produced, suggesting that large differences would be observed for mVOCs based on the culture conditions. To test this idea, we characterized the volatilomes of *Chromobacterium violaceum* (strain ATCC^®^ 12472) and *C. vaccinii* (strain MWU328), and those of their isogenic *cviR*– quorum sensing mutants cultured on solid versus liquid King’s Medium B media. VOCs were sampled using thin-film solid-phase microextraction (TF-SPME) and analyzed by two-dimensional gas chromatography–time-of-flight mass spectrometry (GC×GC-TOFMS). Of the three variables examined – *Chromobacterium* species, media type, and quorum sensing ability – growth on liquid versus solid media caused the most significant differences in the volatilomes. Bacterial species and quorum sensing ability were also influential, but to a lesser degree. Our findings indicate the importance of growth conditions in microbial volatilomics, and therefore, more consideration should be given to how microorganisms are cultured for volatilome analyses.

**Importance:** The purpose of this work is to elucidate the differences in the volatile metabolic profiles of *Chromobacterium spp*. by exploring them through the lens of three variables: growth conditions, species, and the ability to quorum sense. Work on organismal metabolic differences stemming from factors such as liquid versus solid media types remains broadly overlooked. Understanding these effects will allow future researchers to design more robust experiments that better translate to native microbial ecosystems such as rhizosphere and phyllosphere, where volatile compounds may influence plant-pathogen or plant-saprobe interactions.

## Introduction

The health of a soil ecosystem depends on the interactions of the microbes living within it (1, 2). Plants and their root systems, as well as prokaryotic and eukaryotic microorganisms in soils utilize soluble, semi-volatile, and volatile metabolites for communication and regulation of phenotypes (e.g., biofilm formation, motility, virulence factors) within these microbial communities (3-9). Volatile organic compounds (VOCs) play an important role as signaling compounds because their high vapor pressure and low boiling points allow them to traverse discontinuous systems and facilitate longer distance organismal interactions (10, 11). Bioactive VOCs facilitate inter- and intra-organism communication and community dynamics (12-16), where interactions may be beneficial (17, 18) or detrimental to the other organisms (19-23), and VOCs produced by *Bacillus spp sensu lato, Staphylococcus spp, Pseudomonas spp, Rhizobacter spp*, and *Chromobacterium* species are being investigated as potential biocontrol agents against common agricultural pathogens (23-26).

*Chromobacterium* is a genus of gram-negative bacteria typically found in soil and water within tropical and sub-tropical regions (27). Most research on *Chromobacterium* spp. has focused on the characteristic production of violacein, a purple non-diffusible pigment that has antimicrobial properties and potential applications to biotechnology (28). Violacein biosynthesis is regulated through the CviI/CviR quorum sensing (QS) system (29, 30), and in QS-deficient *Chromobacterium* isolates, solid agar and liquid media cultures are unpigmented due to the lack of violacein production (30, 31). This unique visual indication of quorum-dependent expression makes *Chromobacterium* a useful QS reporter strain as well as a common subject of QS research, including the involvement of QS in regulating bioactive VOCs (23, 32-34).

*Chromobacterium* spp. exhibit a range of biological activity, most notably insecticidal activity in a species-specific manner (35-40). Like many other bacteria and fungi, *Chromobacterium* spp produce rich and varied volatilomes of hundreds of microbial VOCs (mVOCs) (17, 23, 41), most of which are yet to be identified or characterized for bioactivity. To identify novel and potentially useful secondary metabolites from a single microbial source, the One Strain-Many Compounds (OSMAC) approach is commonly utilized, in which microbes are cultured under a variety of conditions (e.g., pH, oxygen availability, nutrient availability) to elicit the production of condition-specific compounds (42-44). This strategy has helped to deepen our knowledge of the genetic and regulatory potential for the production of mVOCs (17, 45-52). However, changing the culture conditions from liquid to solid media is rarely included in the OSMAC protocol, though a study of *Aspergillus terreus* showed this variable to have the largest impact on its soluble metabolome among the variables that were tested (44). In previous work, we demonstrated that QS-regulated VOCs of *Chromobacterium vaccinii* cultured on King’s Medium B (KMB) agar inhibit the growth of plant-associated fungi (23), but in unpublished experiments, we observed that the VOCs of *C. vaccinii* cultured in KMB broth were unable to inhibit fungal growth; thus, we posited that solid vs. liquid culture conditions had a significant influence on the volatilomes of *Chromobacterium* spp. In this study, we examined the influence of solid vs. liquid media on the volatilomes of two *Chromobacterium* species, *C. violaceum* and *C. vaccinii*, and their respective isogenic *cviR*– QS mutants.

## Results

### *In vitro Chromobacterium* volatilome

We cultured two quorum sensing sufficient (QS+) strains from *C. violaceum* strain ATCC^®^ 12472 and *C. vaccinii* strain MWU328 and their isogenic *cviR*– quorum sensing mutants (QS– strains MWU12472w and MWU328w, respectively) in liquid KMB broth or on solid KMB agar. Six biological replicates of each of the eight sample types (i.e., four strains × two media) were grown until the QS+ isolates were purple (indicating quorum was reached), at which time the headspace VOCs were collected from all samples using thin film microextraction (TFME). We characterized the volatilomes using comprehensive two-dimensional gas chromatography–time-of-flight mass spectrometry (GC×GC–TOFMS), yielding 189 VOCs after data preprocessing and filtering (Table S1; Fig. S1). Of these, 42 VOCs were chemically identified at level 1 or 2 according to the Metabolomics Standards Initiative (MSI) reporting standards (Table 1) (53). An additional 16 VOCs were identified as level 3 and are reported by their chemical classes, and the remaining 131 VOCs were assigned as level 4 compounds and are reported as unknowns.

**TABLE 1:** MSI level 1 and 2 *Chromobacterium* VOCs and the sample types they were detected in, labeled as liquid vs. solid media, *C. violaceum* vs. *C. vaccinii* (*vio* vs. *vac*), and quorum sensing sufficient vs. *cviR*– quorum sensing mutant strains (QS+ vs. QS–). An open circle indicates that the VOC was detected in at least one of the six biological replicates per sample type, while a solid circle indicates that the VOC was detected in a majority (4 or more) of the six biological replicates.

| Compound Name | ID # | ID Level | Liquid *vac* QS+ | Liquid *vac* QS– | Liquid *vio* QS+ | Liquid *vio* QS– | Solid *vac* QS+ | Solid *vac* QS– | Solid *vio* QS+ | Solid *vio* QS– |
| --- | --- | --- | --- | --- | --- | --- | --- | --- | --- | --- |
| 3-Methylbutan-2-one | KET-242 | 2 | ○ | ● | ● | ● | ○ | ○ | ● | ○ |
| 3-Methylhexan-2-one | KET-554 | 2 | ● | ● | ● | ● | ○ | ○ |  | ○ |
| 3-Methylpentan-2-one | KET-392 | 2 | ● | ● | ○ | ○ |  |  | ○ | ○ |
| Butan-2-one | KET-145 | 1 | ● | ● | ● | ● | ● | ● | ● | ● |
| Decan-2-one | KET-1699 | 1 | ● | ● | ● | ● | ● | ● | ● | ● |
| Dodecan-2-one | KET-2127 | 1 | ● | ● | ● | ● | ● | ● | ● | ● |
| Heptan-2-one | KET-706 | 1 | ● | ● | ● | ● | ● | ● | ● | ● |
| Heptan-3-one | KET-686 | 2 | ● | ● | ● | ● | ● | ● | ● | ● |
| Heptan-4-one | KET-650 | 2 | ● | ● | ● | ● | ● | ● | ● | ● |
| Hexan-3-one | KET-373 | 2 | ● | ● | ● | ● | ● | ○ |  | ○ |
| Nonan-2-one | KET-1388 | 1 | ● | ● | ● | ● | ● | ● | ● | ● |
| Octan-3-one | KET-1005 | 2 | ● | ● | ● | ● | ● | ● | ● | ● |
| Propan-2-one | KET-42 | 1 | ● | ● | ● | ● | ● | ● | ● | ● |
| Tridecan-2-one | KET-2269 | 1 | ○ | ● | ● | ● | ○ | ● | ● | ● |
| Undecan-2-one | KET-1929 | 1 | ○ | ● | ● | ● | ○ | ● | ● | ○ |
| 2-Methylbut-3-en-2-ol | ALC-165 | 2 | ● | ● | ● | ● | ● | ● | ● | ● |
| 3-Methylbut-2-en-1-ol | ALC-457 | 2 | ● | ● | ● | ● | ● | ● | ● | ● |
| 3-Methylbut-3-en-1-ol | ALC-372 | 2 | ● | ● | ● | ● | ● | ● | ● | ● |
| 3-Methylbutan-1-ol | ALC-379 | 2 | ● | ● | ● | ● | ● | ● | ● | ● |
| Butan-1-ol | ALC-261 | 1 | ● | ● | ● | ● | ● | ● | ● | ● |
| Decan-1-ol | ALC-1881 | 1 | ● | ● | ● | ● | ○ | ○ | ● | ● |
| Ethanol | ALC-19 | 1 | ● | ○ | ○ | ○ | ● | ● | ● | ● |
| Hexan-1-ol | ALC-679 | 1 | ● | ● | ● | ● | ● | ● | ● | ● |
| Nonan-1-ol | ALC-1642 | 1 | ● | ○ | ● | ○ | ● | ● | ● | ● |
| Nonan-2-ol | ALC-1411 | 1 | ● | ● | ○ | ○ | ● | ● | ○ | ○ |
| Octan-1-ol | ALC-1333 | 1 | ● | ● | ● | ● | ● | ● | ● | ● |
| Pentan-1-ol | ALC-437 | 1 | ● | ● | ● | ● | ● | ● | ● | ● |
| Propan-1-ol | ALC-111 | 1 | ● | ● | ● | ● | ● | ● | ● | ● |
| 2-Methylpropyl acetate | EST-397 | 2 |  | ○ | ● | ○ |  |  |  |  |
| Ethyl acetate | EST-149 | 1 | ● | ● | ● | ● | ● | ● | ● | ● |
| Propyl acetate | EST-302 | 2 | ● | ● | ● | ● | ● | ● | ● | ● |
| 2-Ethyl-5-methylpyrazine | ARO-1046 | 2 | ● | ● | ○ | ● | ○ | ● | ○ | ● |
| 4-Methylquinazoline | ARO-2119 | 2 | ● | ● | ○ | ● | ○ |  | ○ | ○ |
| Chloromethylbenzene | ARO-1162 | 2 | ● | ○ | ○ | ○ |  |  | ○ | ○ |
| Pyrazine | ARO-339 | 2 | ● | ● | ● | ● | ● | ● | ● | ● |
| Toluene | ARO-388 | 1 | ● | ● | ● | ● | ● | ● | ● | ● |
| 4-Methyl-1-(1-methylethyl)<br>bicyclo[3.1.0]hex-2-ene | HYD-1020 | 2 | ● | ● | ○ | ● | ○ |  | ○ |  |
| Tridecane | HYD-1836 | 1 | ● | ● | ● | ● | ● | ● | ● | ○ |
| Undecane | HYD-1263 | 1 | ○ | ● | ○ | ○ |  |  | ○ |  |
| Methyl thiolacetate | SUL-281 | 2 | ● | ● | ● | ● | ● | ● | ● | ● |
| (E)-But-2-enal | ALD-244 | 2 | ● | ● | ● | ● | ● | ● | ● | ● |
| (E,E)-2,4-Hexadienal | ALD-836 | 2 | ○ | ○ | ○ |  | ○ |  |  | ● |

Pooling the data across *in vitro* sample types, we explored the chemical composition of the *Chromobacterium* volatilome. Only level 1, 2, and 3 VOCs (n = 58) were considered and sorted into seven named chemical classes that represented 97% of the level 1-3 VOCs by counts and 99% of the VOCs by peak abundances, with an undefined chemical class (“Other”) representing the remaining 2% and <1%, respectively. We found that the VOC counts in each chemical class did not follow the same patterns as the VOC abundances. The aldehyde, hydrocarbon, and ester classes made up a lower proportion of the total area compared to the number of VOCs detected (Fig. 1). Conversely, compounds in the sulfur-containing, alcohol, and ketone classes made up larger proportions of the total relative abundance than the number of VOCs detected, with the ketone and alcohol classes representing almost three-quarters of the *Chromobacterium* volatilome by relative abundance. Notably, the sulfur-containing class is represented by just three compounds, methyl thioacetate (SUL-281), dimethyl disulfide (SUL-352), and dimethyl trisulfide (SUL-979), yet makes up 17% of the characterized volatilome by relative abundance.

**FIG 1:**
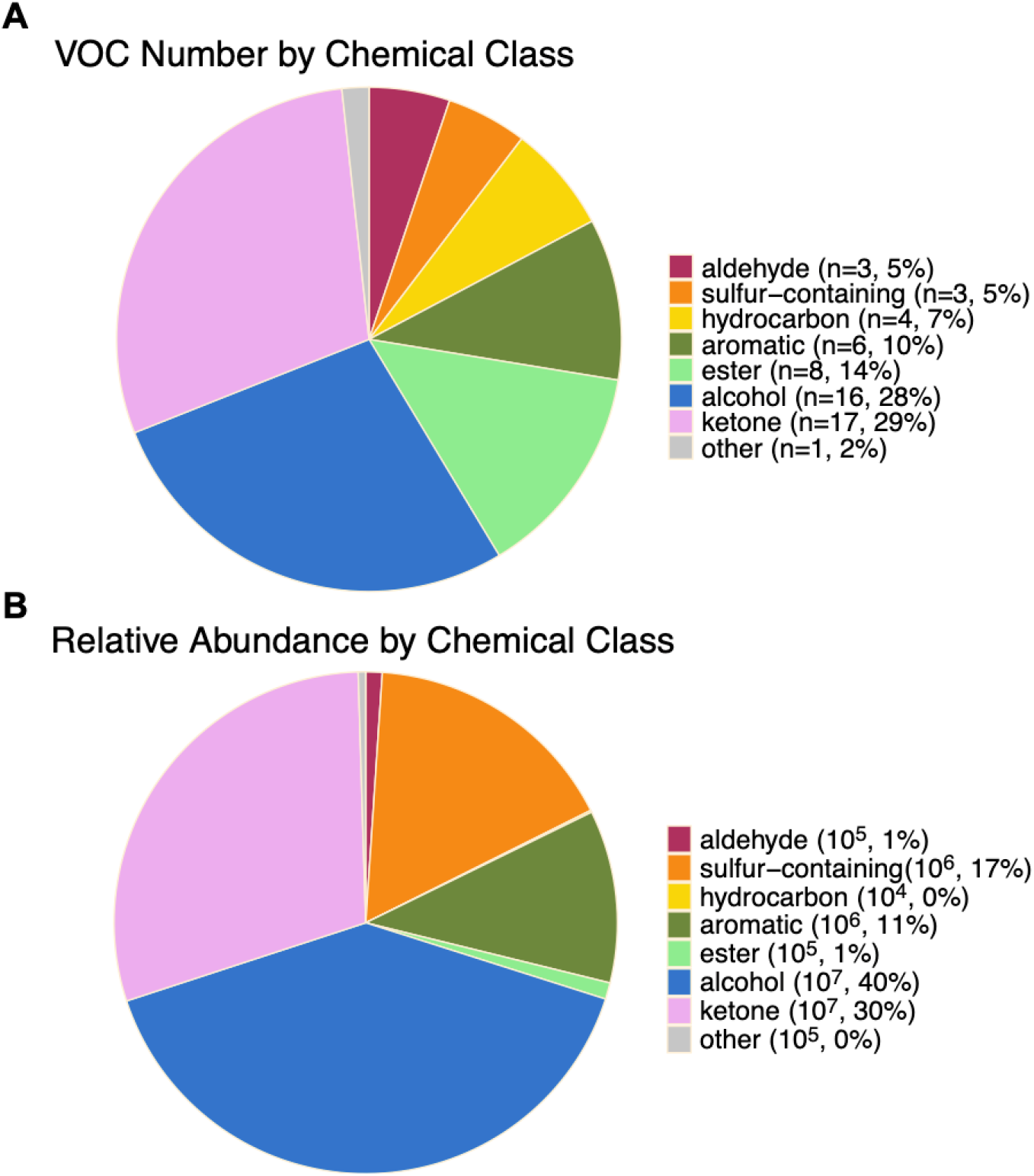
Chemical composition of the *Chromobacterium* volatilome across all sample types and replicates, represented by the (A) total number of VOCs and (B) relative abundance of VOCs, for each chemical class. MSI level 1, 2, and 3 compounds (n = 58) were included in this analysis. In parentheses, the VOC count or average peak area (reported as an order of magnitude, in arbitrary units) is shown, with the percentage indicating the proportion of the total.

### Impact of abiotic and biotic variables on the *Chromobacterium* volatilome

Less than one-third of the *Chromobacterium* VOCs (55 of 189 VOCs) were detected in all eight sample types (Fig. 2, bar 8A), prompting us to explore the roles of biotic (species, QS+/–) and abiotic (media) variables on the volatilome through three binary comparisons: *C. violaceum* vs. *C. vaccinii* (*vio* vs. *vac*), QS sufficient vs. *cviR*− QS mutants (QS+ vs. QS–), and liquid vs. solid media growth conditions. By the presence vs. absence of VOCs, we found that solid vs. liquid growth conditions had the strongest effect on the volatilomes, while strain differences had the weakest effect, with 39% of the volatilome being unique to solid vs. liquid growth, 24% to *vio* vs. *vac*, and 21% to QS+ vs. QS– strains (Fig. S2).

**FIG 2:**
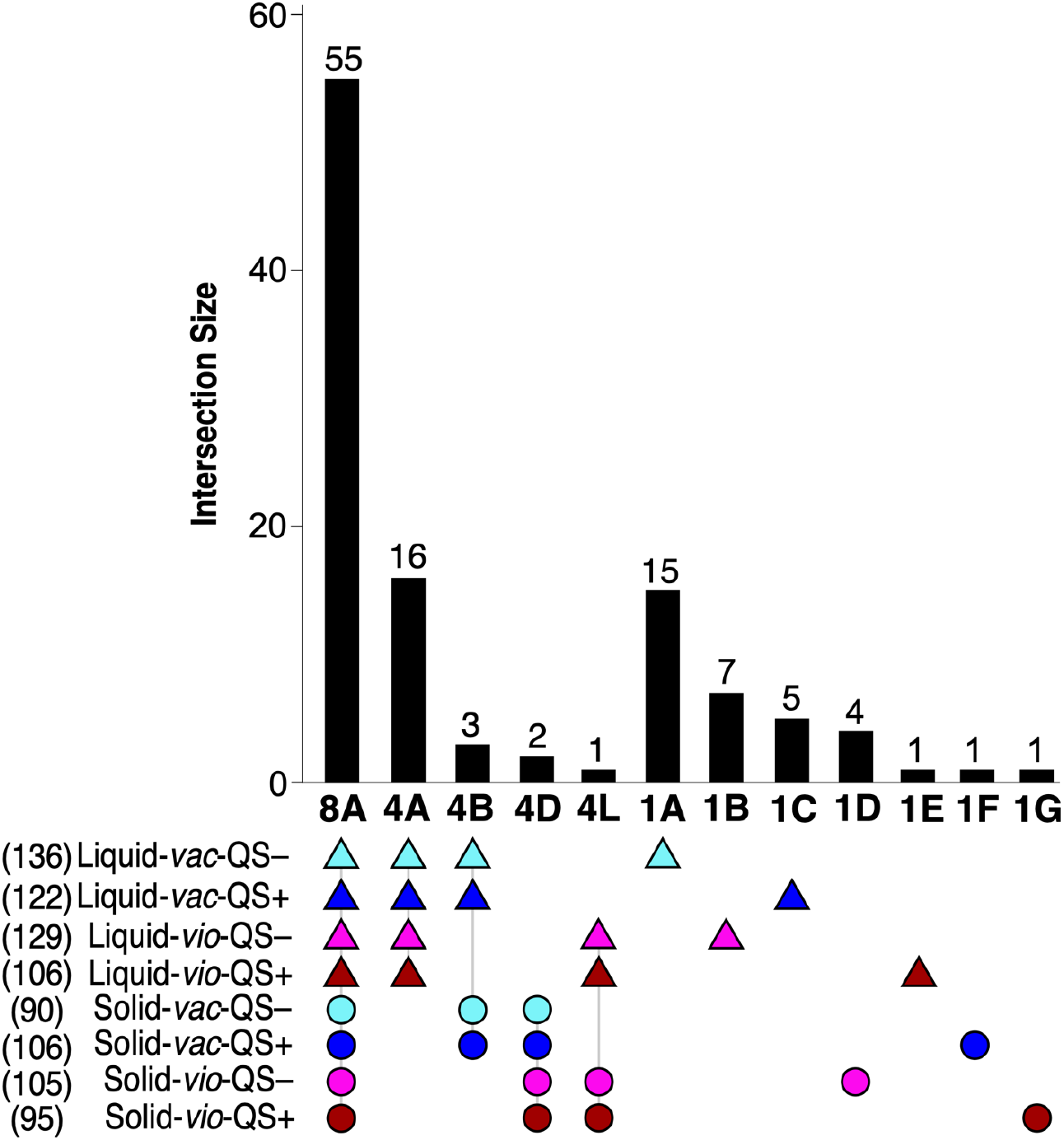
UpSet plot of the *Chromobacterium* volatilome depicting the number of VOCs shared among the eight different sample types. Color and shape indicate whether *C. violaceum* QS+ ATCC^®^ 12472 (red), *C. violaceum* QS– mutant MWU12472w (pink), *C. vaccinii* QS+ MWU328 (blue), or *C. vaccinii* QS– mutant MWU328w (cyan) were grown on solid KMB agar (circle) or liquid KMB broth (triangle). The total number of VOCs in each sample type are in parentheses. The numbers on top and bottom of each vertical bar correspond to the number of VOCs shared between the indicated sample types and the number of sample types in that grouping, respectively. For example, bar 4A indicates that 16 of the 189 VOCs were detected in four sample types. These data are excerpted from Fig. S3.

Of the 54 VOCs unique to liquid samples (Fig. S2), 16 were detected in all liquid samples (30%; Fig. 2, bar 4A), whereas of the 19 VOCs unique to solid samples (Fig. S2), only two were detected in all solid samples (10%; Fig. 2, bar 4D). Very few VOCs were unique to each species while also being conserved across both growth conditions and strains; *C. vaccinii* produced 30 unique VOCs and *C. violaceum* produced 16 (Fig. S2), but only three of the *C. vaccinii* VOCs are detected in both solid and liquid media, while only one *C. violaceum* VOC fits that pattern (Fig. 2, bars 4B and 4L, respectively). There were no VOCs unique to *Chromobacterium* QS+ or to *Chromobacterium* QS– that were conserved across both media and species (Fig. S2). We found that 34 VOCs (18% of the 189 detected) were detected in just one sample type (Fig. 2, bars 1A–1G), the plurality of which were produced by the liquid-*vac*-QS– samples (Fig 2., bar 1A), whereas the solid*-vac-*QS– samples did not produce any unique VOCs.

Overall, significantly more VOCs were detected in liquid samples (n = 170) than in solid samples (n = 135; p < 0.001), but the relative proportions of the seven chemical classes did not significantly differ between sample types (Fig. 3A and B). When considering the relative abundance of the VOCs, differences in the volatilomes were observed (Fig. 3C and D). For example, though most of the seven aromatic VOCs were detected in both liquid and solid samples, the relative abundance of aromatic compounds was significantly higher in liquid (p < 0.001). Most notably, solid-*vac-*QS+ samples exhibited a unique volatilome with a seven-fold increase in the relative abundance of sulfur-containing compounds relative to the other sample groups, almost entirely due to an increase in methyl thioacetate (SUL-281). There was also a two-fold decrease in the abundance of alcohols in the solid-*vac*-QS+ samples relative to the others; however, this was attributable to decreases in several VOCs rather than to a single compound.

**FIG 3:**
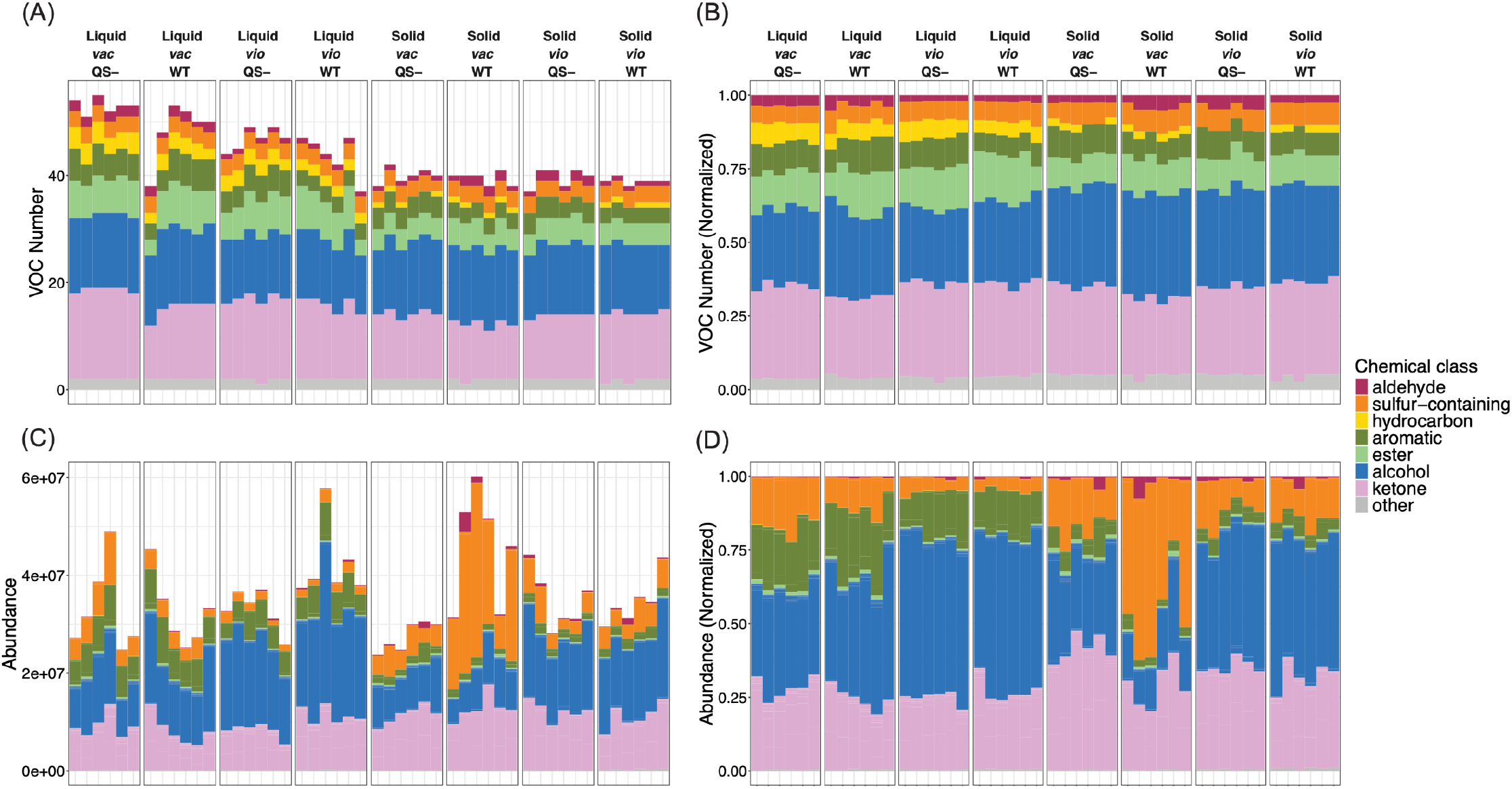
Composition of the level 1 – 3 VOCs of the *Chromobacterium* volatilome by chemical class for each sample type, with each stacked bar representing one biological replicate. The total number of VOCs detected by chemical class, not normalized (A) and normalized (B). The relative abundances of the VOCs detected by chemical class, not normalized (C) and normalized (D).

**FIG 4:**
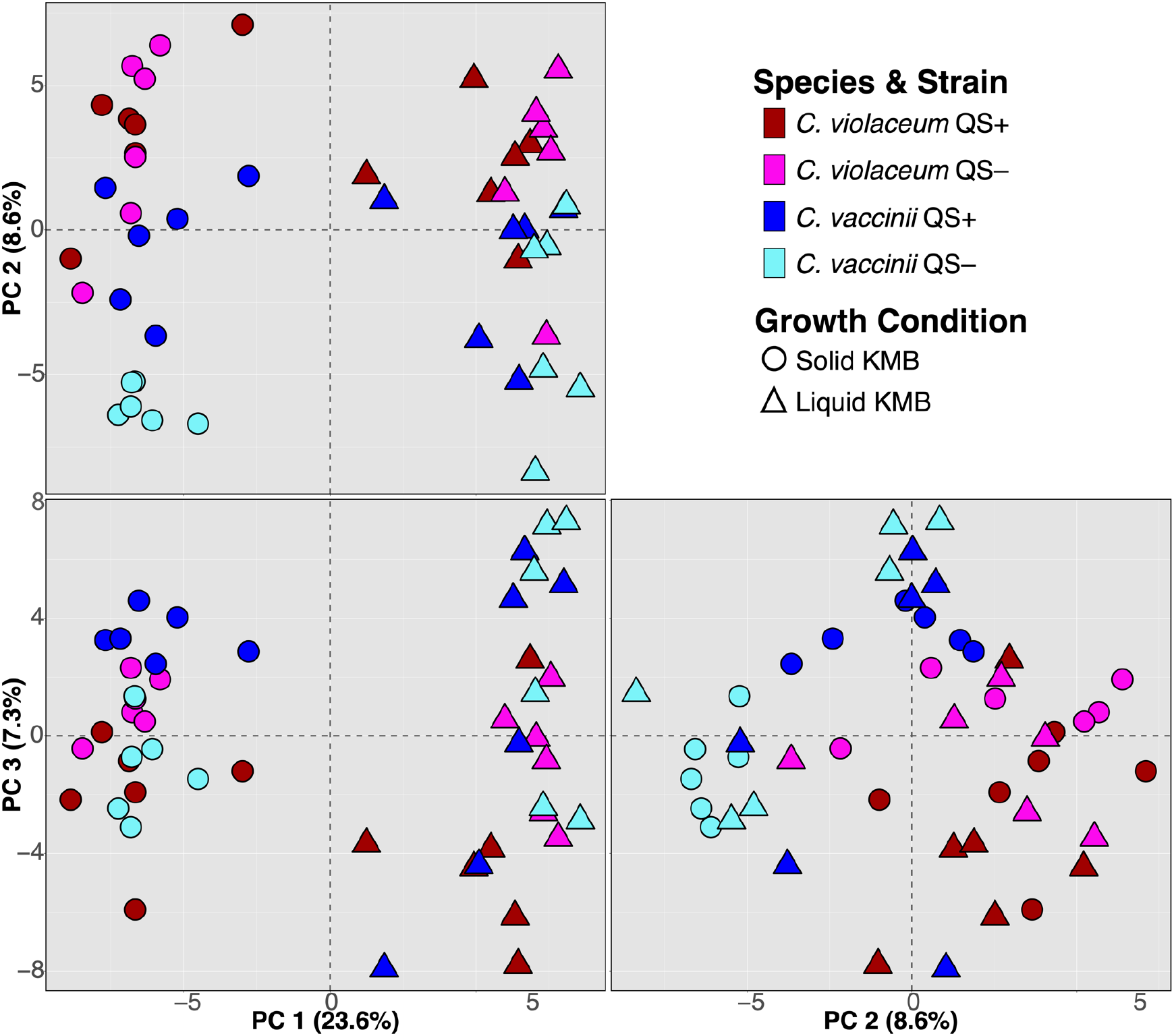
Principal component analysis (PCA) score plot of *Chromobacterium* cultures as observations and 189 VOCs as variables. Six biological replicates were performed for each sample type: *C. violaceum* QS+ ATCC^®^ 12472 (red), *C. violaceum* QS*−* 12472w mutant (pink), *C. vaccinii* QS+ MWU328 (blue), and *C. vaccinii* QS– 328w mutant (cyan) on solid (circles) and liquid (triangles) media.

We performed principal component analysis (PCA; Fig. 4) and hierarchical clustering analysis (HCA; Fig. 5) as an unsupervised approach to explore the relationships between each of the *Chromobacterium* sample types based on the VOCs that were detected. Using the six biological replicates of each culture as observations and the 189 *Chromobacterium* VOCs as variables, the PCA revealed that the largest proportion of variance in the model (PC1, 23.6%) was explained by liquid vs. solid growth conditions (Fig. 4); liquid cultures of both species (i.e., *C. violaceum* and *C. vaccinii*) clustered together along PC1 > 0, while solid cultures of both species clustered away from liquid cultures along PC1 < 0. The distinct clusters of solid vs. liquid replicates along PC1 were congruent with the differences we observed in the numbers of VOCs detected (Fig. S2) and the chemical class abundances between the respective volatilomes (Fig. 3). Additionally, the volatile profiles of the samples were observed to separate based on *Chromobacterium* species in PC2 vs. PC3. The influence of QS ability on the *Chromobacterium* volatilome was relatively small in comparison to that of the growth condition and species, as reflected by the lack of clear separation between the QS+ and QS– samples within the PCA. Representing the *Chromobacterium* volatilome in PCA biplots (Fig. S4) revealed that many of the compounds unique to all liquid samples (Fig. 2, bar 4A) and all solid samples (Fig. 2, bar 4D) had strong loadings on PC1, contributing to the separation of liquid and solid samples.

**FIG 5:**
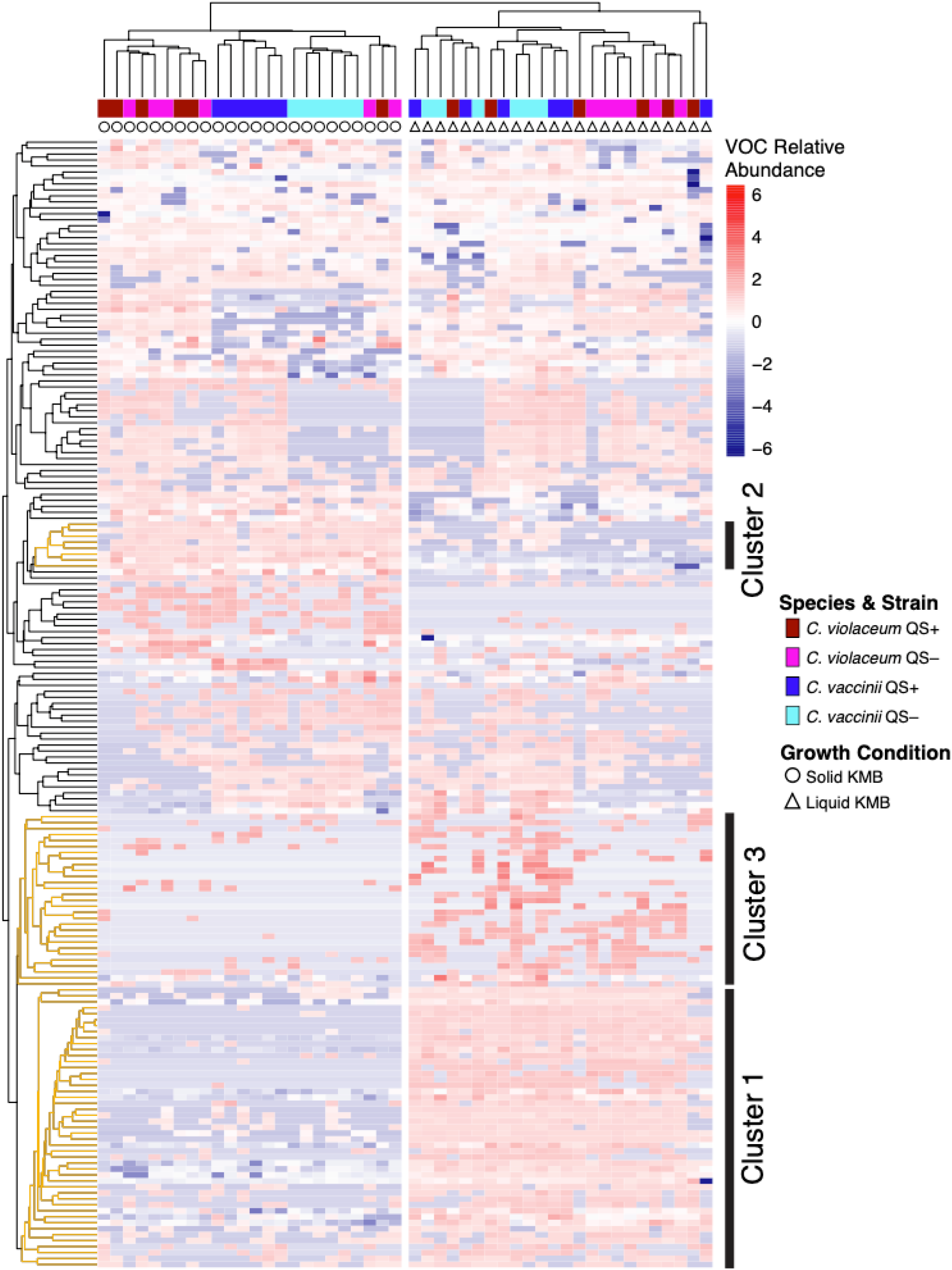
Hierarchical clustering analysis (HCA) of the 48 biological replicates (columns) and the 189 VOCs (rows), using Manhattan distance and average linkage. The relative abundance of each VOC is depicted by the color gradient from blue (low abundance) to red (high abundance). The sample type for each biological replicate is indicated by the marker shape (growth condition) and color (species and QS) as labeled in the key. Gold-colored branches of the rows correspond to VOC Clusters 1, 2, and 3.

Similar to the observations from the PCA, clustering of samples in the HCA occurred primarily according to growth conditions (Fig. 5). Along the columns of the HCA, biological replicates clustered primarily by solid vs. liquid media, and we identified two major clusters of VOCs that were associated with liquid vs. solid growth. Cluster 1 VOCs, containing 47 compounds, were more abundant in the headspace of liquid media samples and included all 16 VOCs that are unique to all liquid samples (Fig. 2, bar 4A). Cluster 2 VOCs, containing eight compounds, were more abundant in solid samples and included the two VOCs uniquely produced by all solid samples (Fig 2A, bar 4D). Additionally, many of the compounds unique to one class of liquid samples (Fig 2, bars 1C and 1E) are found in a third cluster comprised of 29 VOCs. Within the two primary sample clusters separated by growth condition, there is additional clustering by *Chromobacterium* species. Notably, the biological replicates of *C. vaccinii* grown on solid media clustered in two groups separated by QS+ vs. QS–, whereas samples of *C. violaceum* grown on solid media and cultured in liquid media did not.

## Discussion

We previously characterized the production of fungistatic VOCs by *C. vaccinii* strain MWU328 (23). However, during method development for that study, we observed that the production of fungistatic VOCs was context dependent; the VOCs of *C. vaccinii* cultured on solid KMB agar were strongly fungistatic, but VOCs from liquid cultures of *C. vaccinii* were not (unpublished). That phenomenon motivated this work, to investigate the influence of solid vs. liquid media on the volatilomes of quorum sensing-sufficient and quorum sensing-deficient mutants of *Chromobacterium* spp. Our data show that changing the growth conditions had a large impact on the volatile profiles of the isolates we studied, which was mostly attributable to VOCs that are uniquely produced in one growth condition or the other, but differences in the relative abundances of VOCs produced in both solid and liquid were also observed. These findings are consistent with prior studies focused on differences in soluble metabolites based on growth conditions (44-46, 54-62). For example, Alhameed et al. explored the profiles of *Candida albicans* cultured in yeast potato dextrose media and found that most soluble metabolites were significantly greater in abundance in solid cultures (58), and Fan et al. noted little overlap in the soluble metabolite profiles of ten marine-derived fungal strains when comparing liquid and solid cultures (56).

Of the 189 VOCs detected in this study, we were able to assign putative names to 42 compounds, 40 of which are known mVOCs (41), and six have been previously reported as part of the *Chromobacterium* volatilome: hexan-1-ol, octan-1-ol, 3-methylbuten-1-ol, nonan-2-one, undecan-2-one, 3-hydroxybutan-2-one (17, 23). Undecan-2-one was detected in five of the eight sample types (Fig. S3, bar 5I), and the other five previously reported VOCs were detected in all eight sample types (Fig 2, bar 8A). Additionally, 24 of the named VOCs that we detected have known bioactive properties, including but not limited to antifungal activity (41). For example, 3-methyl-pentan-2-one, detected here in *C. violaceum* liquid samples, has been demonstrated to have fungistatic activity against *Paecilomyces lilacinus* and *Clonostachys rosea* (63), and undecan-2-one has been shown to exhibit antifungal activity against *Rhizoctonia solani* (64), *Sclerotinia sclerotiorum* (26, 64, 65), *Fusarium oxysporum* (64, 66), and *Alternaria radicina* (64). Many bacteria and fungi utilize the production of sulfur-containing compounds as a biocontrol mechanism (67-74), and the three sulfur-containing compounds that were detected in this study—methyl thioacetate, dimethyl disulfide, and dimethyl trisulfide—have previously been reported as biocontrol compounds against phytopathogenic nematodes, bacteria, and fungi (75-79), and methyl thioacetate as a promoter of tomato plant growth (80). These three compounds were detected in all sample types, but methyl thioacetate was significantly more abundant in *C. vaccinii* QS+ MWU328 cultured on solid media. This compound may, therefore, be responsible entirely or in part for the QS-dependent inhibition of fungal growth that we previously observed for this *C. vaccinii* strain when cultured on KMB agar (23).

It should be noted that this study included only one strain for each *Chromobacterium* species (i.e., *C. violaceum* strain ATCC^®^ 12472 and *C. vaccinii* strain MWU328), which limits our ability to generalize our findings about the influences of growth conditions and QS on the volatilome to other *Chromobacterium* strains and species. As one example, we observed that cultures of *C. vaccinii* strain MWU328 grown on solid media clustered by QS ability (Fig 4; Fig. 5); however, this influence was not observed for *C. violaceum* strain ATCC^®^ 12472, and we do not know if this phenomenon would be observed for other *C. vaccinii* or *C. violaceum* strains.

Though this is the most comprehensive investigation of the *Chromobacterium* volatilome to date, much more work is required for a full characterization. Prior studies have demonstrated a significant amount of intra-species variation in microbial volatile profiles (47, 81-86), and therefore, we expect that we would need to characterize dozens more strains of *C. violaceum* and

*C. vaccinii* to estimate the core and accessory volatilomes of these species (81, 85), and this effort would need to be repeated for the other species of *Chromobacterium* to characterize the volatilome of the genus; there are 19 currently recognized species of *Chromobacterium* plus others yet to be recognized.

Often, microbial volatilomes are studied to identify VOCs of ecological importance, whether the microbes dwell in environmental reservoirs, animal hosts, or built environments. However, due to the biological complexity of those microbial ecosystems and the technical complexity of sampling mVOCs from their native environments, many studies are initially carried out using *in vitro* monocultures, and the biological relevance of those volatilomes are later explored. The vast majority of the mVOC literature is from studies that used liquid cultures, or the culture conditions are not specified (41). Given the significant soluble and volatile metabolomic reprogramming that occurs when bacteria and fungi switch from planktonic to sessile growth (47, 56, 58, 87, 88), matching the *in vitro* growth conditions to the microbial growth conditions of the native habitat is likely to yield more translatable results. At a minimum, efforts need to be made to more thoroughly document all variables of an experimental setup to facilitate volatilome comparisons between studies.

## Conclusions

Growth within liquid versus solid media was the most important variable in altering the volatilomes of *C. violaceum* and *C. vaccinii*. We identified significant differences in the relative abundances of alcohols and sulfur-containing compounds between growth conditions and an overall greater number of VOCs detected in liquid samples under the conditions we tested. Our results show that the production of *Chromobacterium* VOCs, including known bioactive VOCs, can be appreciably affected by planktonic versus sessile growth and highlight the importance of studying growth on solid media for the robust characterization of in vitro volatile metabolomes. There is a paucity of information, however, regarding the volatilomes of microbes cultured on solid media; consequently, more work is necessary for the robust characterization of microbial volatilomes in vitro.

## Materials and Methods

### Bacterial Isolates and Culture Conditions

The *Chromobacterium* isolates used in this study were quorum sensing sufficient (QS+) strains *C. violaceum* ATCC^®^ 12472 (89) and *C. vaccinii* MWU328 (90), as well as their spontaneous isogenic *cviR*– quorum sensing mutants (QS–) MWU12472w (31) and MWU328w, respectively. The bacteria were cultured in King’s Medium B (KMB) broth and incubated overnight with shaking at 25 ºC. For each of the overnight cultures, 1.2 mL was transferred into 25 mL KMB broth (for liquid culture experiments) and 500 µL plated onto KMB agar (for solid culture experiments). Bacteria cultures were prepared in six biological replicates for each media type (i.e., broth or agar) and corresponding media-only controls were prepared in triplicates. Broth and agar cultures were incubated at 25 ºC for 7 h and 15.5 h, respectively. At these times, violacein production in the wild-type cultures indicated quorum had been reached.

### VOC Collection and Analysis by HS–GC×GC–TOFMS

Headspace VOCs were collected using thin film microextraction (TFME). Upon reaching quorum, three thin films (TFs) composed of divinylbenzene and polydimethylsiloxane (DVB/PDMS) on a carbon mesh were aseptically suspended in the flask headspace for liquid samples or placed on the lid of an inverted Petri dish for solid samples. The cultures and media-only controls, containing TF technical triplicates, were incubated for 1 h at 25 ºC for VOC collection, after which the TFs were removed, sealed in conditioned glass thermal desorption tubes and stored at 4 ºC until volatile metabolomics analysis. VOCs were analyzed using thermal desorption coupled with comprehensive two-dimensional gas chromatography–time-of-flight mass spectrometry (GC×GC–TOFMS). Parameters for thermal desorption, chromatography, and mass spectrometry are in Table S2. An external alkane standards mixture (C_8_–C_20_ in hexane; Sigma-Aldrich, St. Louis, MO, USA) was analyzed for retention index (RI) calculations.

### Data Processing and Compound Identification

Data processing and alignment were performed using Leco ChromaTOF^®^ software with the Statistical Compare package (Table S3). VOCs were identified according to the Metabolomics Standards Initiative (MSI) minimum reporting standards (53). Putative VOC names were assigned by forward matches to the National Institute of Standards and Technology (NIST^®^) 2011 mass spectral library and comparison of experimental retention indices (RIs) to published RIs. Criteria for a level 1 identification were as follows: (i) a mass spectral match and first dimension retention index match (< 6 units difference) with an external standard or (ii) a strong linear relationship (R^2^ ≥ 0.989) of retention times vs. carbon number with external standards in a homologous series (i.e., 1-alcohols, 2-alcohols, and 2-ketones). Criteria for a level 2 identification were as follows: (i) a strong linear relationship (R^2^ ≥ 0.989) of retention times vs. carbon number with compounds in a homologous series without an external standard (i.e., 1-alcohols, 2-ketones, 3-ketones, and 3-methyl-2-ketones) or (ii) (a) ≥ 700 mass spectral match by a forward search of the NIST library and (b) experimental mid-polar 624 RIs consistent with published RIs (i.e., within 12 units) after converting the median NIST^®^-reported non-polar RI to a mid-polar 624 RI (91). Level 3 identifications had (i) ≥ 700 mass spectral match and (ii) an experimental mid-polar 624 RI within 36 units of the median published RI. Level 1, 2, and 3 compounds were assigned to chemical classes based on characteristic chromatographic patterns in second dimension retention times (92-96). Level 3 compounds were reported only as the chemical class name. Level 4 compounds have insufficient mass spectral matches or inconsistent RI data and are reported as unknowns.

### Statistical Analyses

All statistical analyses were performed using R version 4.2.1 (R Foundation for Statistical Computing). Prior to statistical analyses, as summarized in Fig. S1, chromatographic contaminants and artefacts (e.g., siloxanes, atmospheric gases) were removed from the peak table. Missing values for technical triplicates and technical duplicates were imputed using replicate Random Forest or half-minimum values, respectively, from the R MetabImpute package version 0.1.0 (97). The relative abundance of compounds across chromatograms was normalized using probabilistic quotient normalization (PQN) (98). Analytes were retained for further analysis based on the following criteria: (i) significantly greater in abundance in a sample type than corresponding medium (i.e., broth or agar) controls using the Wilcoxon rank-sum test with Benjamini-Hochberg correction and a one-sided alpha of 0.05 and (ii) arithmetic means of technical replicate peak areas were at least two-fold greater in abundance in any biological replicate than the corresponding medium controls. Log_10_ transformation was applied, and the abundances of technical replicates were averaged for each biological replicate for further analyses. The relatedness of biological replicates was explored using principal component analysis (PCA) and hierarchical clustering analysis (HCA) using the samples as the observations and the absolute peak intensities (mean-centered and scaled to unit variance) as the variables. For comparisons of VOC presence or absence between classes, a VOC needed to be detected in at least four out of six biological replicates to have been considered present in the corresponding sample class. The following R packages were used for PCA, HCA, and Upset plot analyses: ggplot2 (3.5.0) (99), pheatmap (1.0.12), UpSetR (1.4.0).

### Data availability

Metabolomic data (chemical feature peak areas and retention time information) included in this study are available at the NIH Common Fund’s National Metabolomics Data Repository (NMDR) website, the Metabolomics Workbench (100), at www.metabolomicsworkbench.org, where it has been assigned project ID PR002570 and study ID ST004091 (https://doi.org/10.21228/M8FN9W).

## Supplemental Materials

**Figure S1**: Flowchart of the steps that were applied to the processed data to prepare it for statistical analyses

**Figure S2**: Venn diagrams comparing the number of VOCs in the *Chromobacterium* spp. volatilome by media type, species, and quorum sensing sufficiency

**Figure S3**: UpSet plot of the *Chromobacterium* spp. volatilome depicting the number VOCs shared among sample types

**Figure S4**: Principal component analysis (PCA) biplot of *Chromobacterium* spp. cultures as observations and 189 VOCs as variables

**Table S1**: Table of all 189 *Chromobacterium* VOCs detected in the headspace of solid and liquid cultures

**Table S2**: Parameters for headspace TFME and GC×GC-TOFMS analysis

**Table S3**: Parameters for data processing and alignment

## Acknowledgements

This project was supported by a National Science Foundation Learning, Entering, Advising, and Producing research (LEAP) scholarship and an Arizona State University School of Life Sciences Undergraduate Research (SOLUR) scholarship to DR.

## Author contributions

HDB conceived of the study; HDB, DR, EAHK, and TJD designed the experiments; DR, EAHK, and TJD collected the data; JAD and JD processed and analyzed the data; JAD, JD, and HDB wrote the manuscript; all authors edited the manuscript and approved the final version.

