## Supplemental Information for "The *Chromobacterium* Volatilome is Strongly Influenced by Growth on Liquid versus Solid Media"

**Table S1: See Supplementary Excel File.** Table of 189 *Chromobacterium* VOCs detected in the headspace of *C. vaccinii* MWU328 and 328w and *C. violaceum* ATCC® 12472 and 12472w cultured on solid and liquid King's Medium Broth (KMB). For each compound the following information is reported: the VOC identification number assigned by the data processing software, chemical abstract service (CAS) identifier number, chemical class, metabolomics standards initiative (MSI) identification level, average first and second dimension retention times ($^{1}t_R$ and $^{2}t_R$, respectively), the observed first-dimension retention index on 624-Sil stationary phase, and if the compound was observed as part of a homologous series. For each of the sample types and media blanks, the number of replicates each compound was detected in is reported along with the mean relative peak areas and standard deviations.

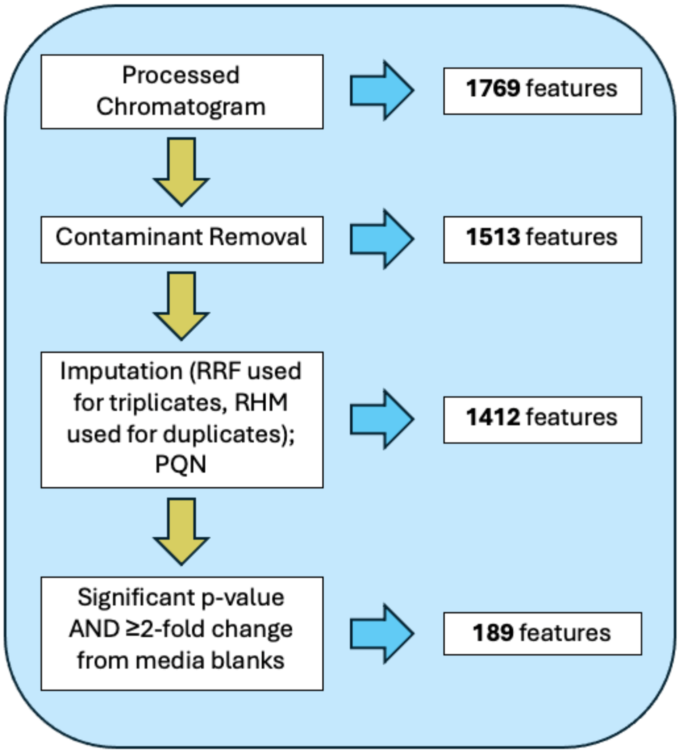


**Figure S1:** Flowchart of the steps that were applied to the processed data to prepare it for statistical analyses. RRF, replicate random forest; RHM, replicate half-minimum; PQN, probabilistic quotient normalization.

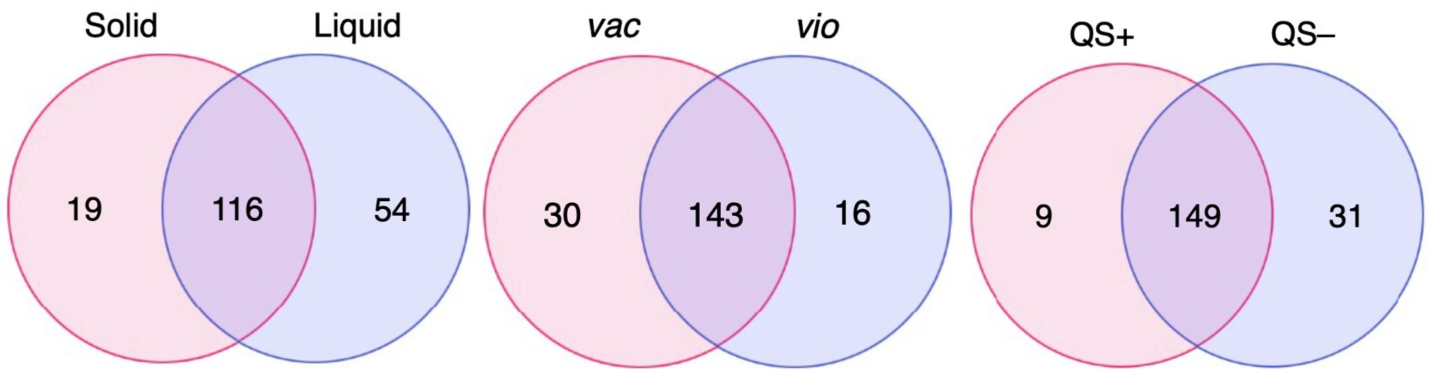


**Figure S2:** Venn diagrams comparing the number of VOCs in the *Chromobacterium* spp. volatilome by media type (solid vs. liquid; left), species (*C. vaccinii* (*vac*) vs. *C. violaceum* (*vio*); center), and quorum sensing sufficiency (QS+ vs. QS–, right).

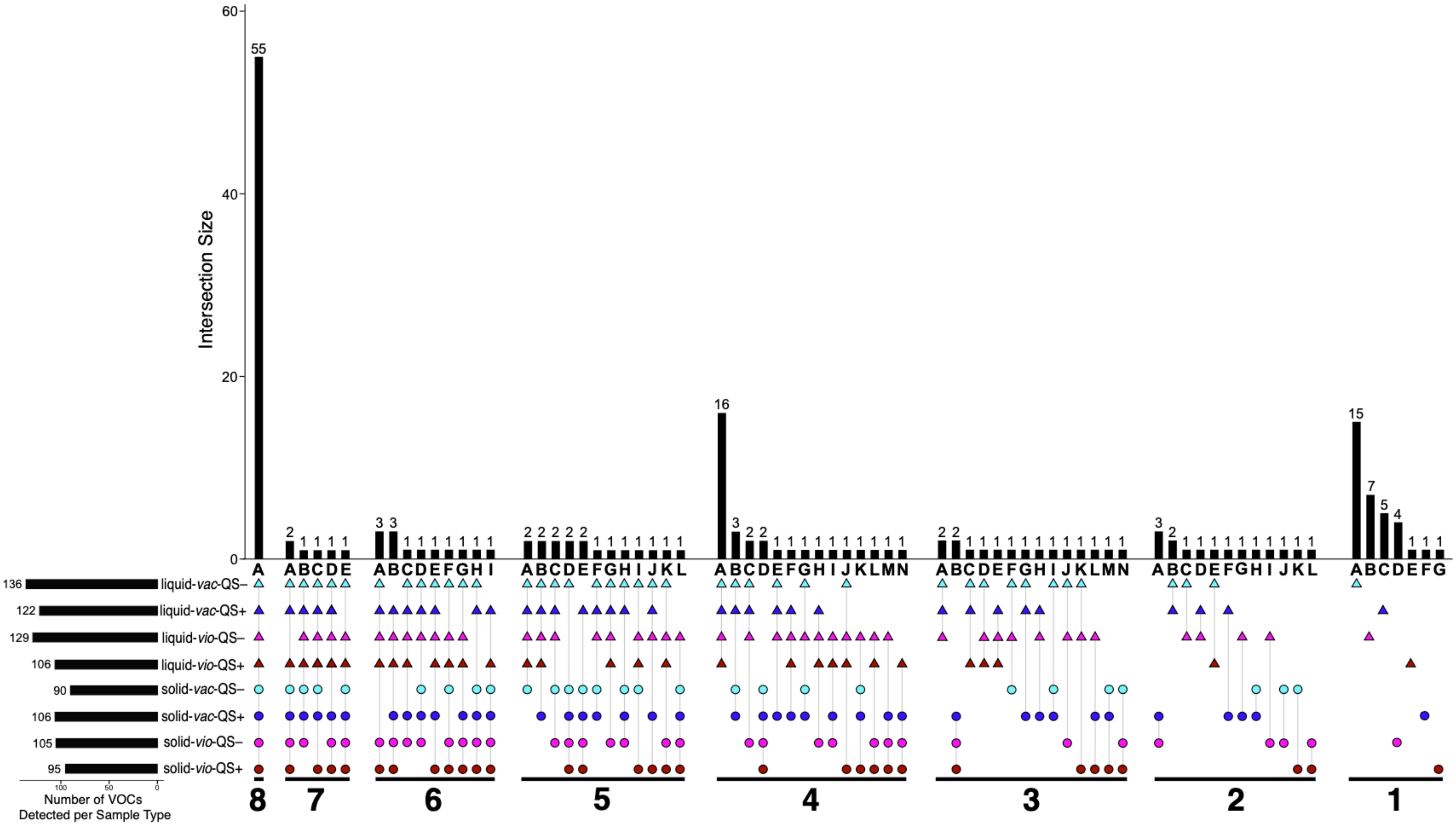
Intersection Size
0
20
40
60
55
16
15
liquid-vac-QS−
liquid-vac-QS+
liquid-vio-QS−
liquid-vio-QS+
solid-vac-QS−
solid-vac-QS+
solid-vio-QS−
solid-vio-QS+
136
122
129
106
90
106
105
95
Number of VOCs
Detected per Sample Type
8
7
6
5
4
3
2
1

**Figure S3:** UpSet plot of the *Chromobacterium* spp. volatilome depicting the number VOCs shared among sample types. Groupings are clustered by number of sample types included (1 – 8) and labeled below each cluster. The number of VOCs shared by each grouping is indicated at the top of each vertical bar. Color and shape indicate *C. violaceum* QS+ ATCC® 12472 (red), *C. violaceum* QS– mutant 12472w (pink), *C. vaccinii* QS+ MWU328 (blue), or *C. vaccinii* QS– mutant 328w (cyan) growth on solid KMB agar (circle) or liquid KMB broth (triangle). The total number of VOCs for each sample type is indicated by the bar graph and corresponding number in each row.

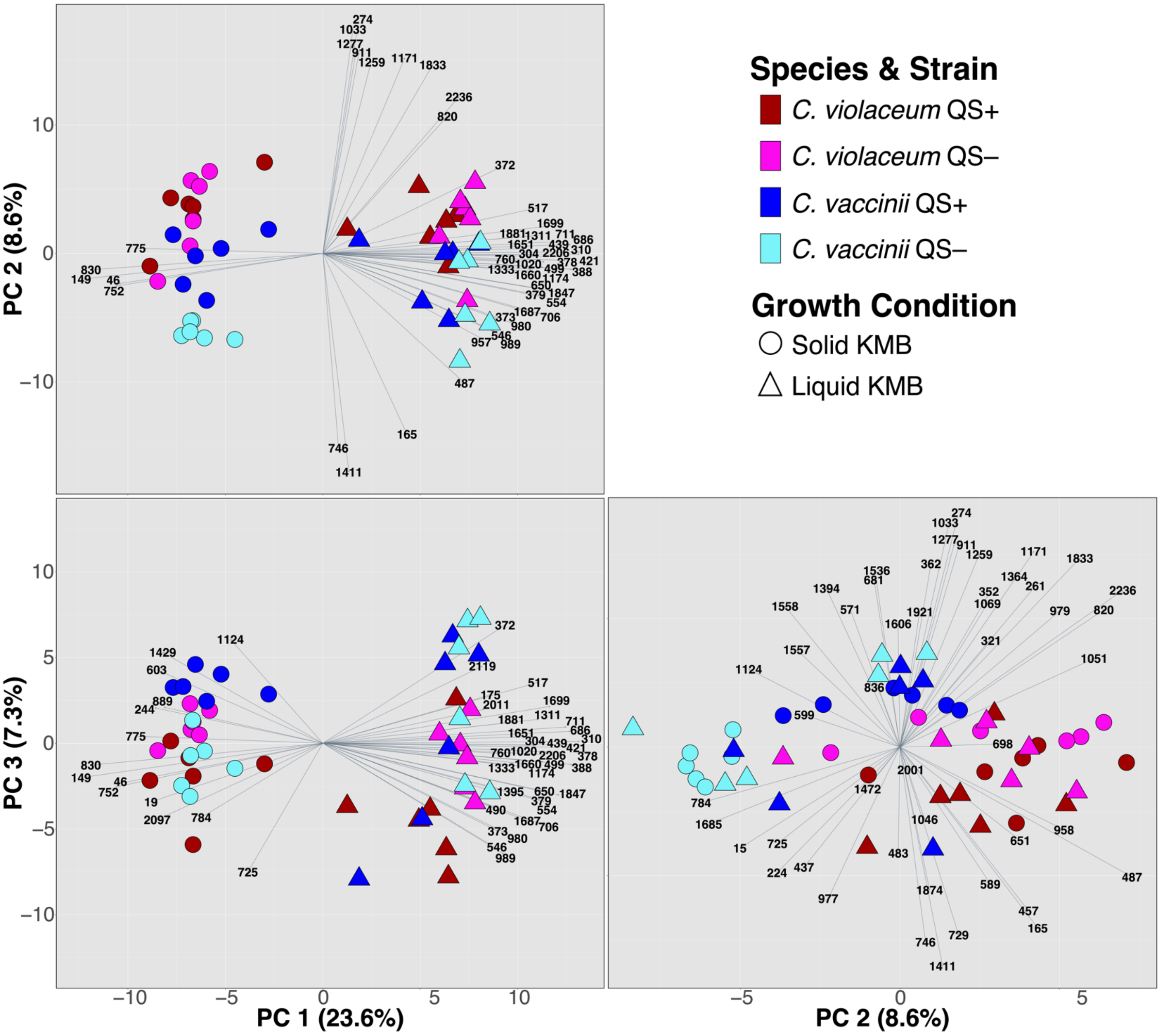


**Figure S4:** Principal component analysis (PCA) biplot of *Chromobacterium* spp. cultures as observations and 189 VOCs as variables. Six biological replicates were performed for each sample type: *C. violaceum* QS+ ATCC® 12472 (red), *C. violaceum* QS– 12472w mutant (pink), *C. vaccinii* QS+ MWU328 (blue), and *C. vaccinii* QS– mutant 328w (cyan) on solid (circles) and liquid (triangles) media. Vectors indicate the top 50 loadings across the principal components. Each vector's number indicates a VOC with the chemical class designation removed (i.e. an arrow with "725" indicates ALC-725 from Table S1).

**Table S2.** Parameters for headspace TFME and GC×GC-TOFMS analysis

| | |
|---|---|
| **AUTOSAMPLER METHOD** | |
| Instrument description | Gerstel® MPS Pro® |
| Software description | Gerstel® Maestro® (version 1.5.3.2) |
| **Thermal Desorption Parameters** | |
| Initial temperature | 50 ºC |
| Ramp rate | 720 ºC·min$^{-1}$ |
| End temperature | 220 ºC |
| Hold time (min) | 3.00 |
| Transfer temperature | 240 ºC |
| Transfer temperature mode | Fixed |
| Desorption mode | Splitless |
| Sample mode | Retain Tube – Standby Cooling |
| Standby temperature | 50 ºC |
| **Inlet (CIS) Parameters** | |
| Initial temperature | -80 ºC (cryo cooling) |
| Equilibrium time | 0.20 min |
| Initial time | 0.0 min |
| Ramp rate | 12 °C·s$^{-1}$ |
| End temperature | 250 °C |
| Hold time | 3.00 min |
| Heater mode | Standard |
| Liner | Glass liner, non-baffled w/glass wool |
| **TWO-DIMENSIONAL GAS CHROMATOGRAPHY METHOD** | |
| Instrument description | Agilent® 7890B |
| Software description | Leco ChromaTOF (version 4.72.0) |
| Column configuration | Column 1: Rxi®-624Sil MS, 60 m × 0.25 mm × 1.4 $\mu$m<br>Column 2: Stabilwax®, 1 m × 0.25 mm × 0.5$\mu$m |
| Carrier gas | Helium, 2 mL·min$^{-1}$ (constant) |
| Front inlet type | Gerstel® |
| Front inlet mode | Splitless |
| Front inlet septum purge flow | 1 mL·min$^{-1}$ |
| Front inlet septum purge time | 300 s |
| Front inlet purge flow | 50 mL·min$^{-1}$ |
| Front inlet total purge flow | 52 mL·min$^{-1}$ |
| Oven equilibration time | 5 s |
| Primary oven temperature ramp | Initial temperature: 35 °C<br>Initial time: 0.5 min<br>Ramp rate: 5 C·min$^{-1}$<br>Final temperature: 230 °C<br>Hold time: 5 min |
| Secondary oven temperature offset | +5 °C (relative to primary oven) |
| Modulator temperature offset | +15 °C (relative to secondary oven) |
| Modulation timing | Modulation period: 2.00 s<br>Hot pulse time: 0.50 s<br>Cold pulse time: 0.50 s |
| Transfer line temperature | 250 °C |

| MASS SPECTROMETRY METHOD | |
|---|---|
| Instrument description | LECO® Pegasus® 4D |
| Use GC method total time for MS method total time | Yes |
| Acquisition delay | 180 s |
| Filament active time | 180 s to end of run |
| Start mass/End mass | 35/400 |
| Acquisition rate | 100 spectra·$s^{-1}$ |
| Optimized voltage offset | +50 V |
| Electron energy | -70 eV |
| Ion source temperature | 250 °C |

**Table S3.** Parameters for data processing and alignment

| DATA PROCESSING METHOD | |
|---|---|
| Software description | LECO® ChromaTOF® with Statistical Compare (version 4.71.0.0) |
| Baseline tracking/Offset | Entire run/0.5 (through middle of noise) |
| Data points averaged for smoothing | Auto |
| First dimension peak width | 12 slices |
| Mass spectral match required to combine | 600 |
| Second dimension peak width | 0.15 |
| Min. subpeak signal-to-noise (S/N) for | 6 |
| Integration approach | Traditional |
| Peak finding | S/N: 50<br>Number of apexing masses: 2 |
| Mass spec libraries for searching | NIST 2011 |
| Library identity search mode | Normal |
| Library search mode | Forward |
| Minimum molecular weight allowed | 35 |
| Maximum molecular weight allowed | 700 |
| Mass threshold (0-998) | 10 |
| Mass to use for area/height calculation | Unique mass |
| Alignment analyte match criteria | Spectral match mass threshold: 10<br>Minimum spectral similarity match: 600<br>Max. number of modulation periods apart: 3<br>Max. retention time difference (s): 0.2<br>S/N for second peak find: 5 |
| Criteria for inclusion of analytes | Min. number of samples that contain analyte: 1 |
